# CSF1R inhibitors pexidartinib and sotuletinib induce rapid glial ablation despite their limited brain penetrability

**DOI:** 10.1101/2025.10.11.681780

**Authors:** Olaf van Tellingen, Zoey J Tolboom, Yoran M Leter, Efi Tsouri, Artur M Burylo, Caroline L van Heijningen, Sanne B Schagen, Mark C de Gooijer, Laura E Kuil

**Author notes:** Corresponding author: Laura E Kuil Division of Psychosocial Research and Epidemiology – NKI-Avl, Plesmanlaan 121, 1066CX Amsterdam, the Netherlands.

## Abstract

**Background:** Microglial reactivity, a hallmark of many neurodegenerative diseases, is thought to contribute significantly to disease pathology. In experimental models, colony stimulating factor 1 receptor (CSF1R) inhibitors transiently deplete microglia to resolve inflammation, leading to improved neuropathology. In oncology, CSF1R inhibitors modulate tumor-associated macrophages (TAMs) toward a tumor-suppressive phenotype by silencing CSF1–CSF1R signaling. As for any therapeutic, target engagement depends on effective drug delivery. In the brain a major hurdle is the limited drug delivery caused by the presence of the blood brain barrier (BBB) containing drug efflux transporters. However, the affinity to these transporters of most CSF1R inhibitors is unknown.

**Methods:** We assessed the brain penetrance of two CSF1R inhibitors, pexidartinib (PLX3397) and sotuletinib (BLZ945), in the absence and presence of drug transporters ABCB1 and ABCG2. We further assessed their impact on peripheral immune populations, tissue-resident macrophages, microglia and oligodendrocyte progenitor cells (OPCs).

**Results:** Both compounds have a limited brain permeability (brain-to-plasma ratio: 0.1). Sotuletinib was a substrate for both ABCB1 and ABCG2, whereas pexidartinib was transported primarily by ABCB1. Despite low brain exposure, both are able to ablate microglia when given to mice at high doses, accompanied by marked depletion of OPCs and macrophage populations in the liver, intestine, and kidney, as well as non-classical monocytes in blood. Pexidartinib additionally altered splenic immune composition, increasing T cells and neutrophils, and reducing dendritic cells and non-classical monocytes.

**Conclusion:** These findings highlight that high-dose CSF1R inhibition rapidly depletes microglia, but induces substantial off-target effects. Such systemic impacts, as well as the impact on OPCs, should be considered when interpreting experimental outcomes or translating CSF1R inhibition into clinical contexts where brain targeting is required.

**Key messages:**

- CSF1R inhibitors pexidartinib and sotuletinib show poor brain penetrance (brain-to-plasma ratio 0.1)
- Sotuletinib is a substrate to ABCB1 and ABCG2, pexidartinib is a substrate to ABCB1.
- Both drugs rapidly deplete microglia, despite poor brain penetration
- Microglia depletion is accompanied by loss of OPCs and tissue macrophages

## Introduction

Colony-stimulating factor-1 receptor (CSF1R) inhibitors attract interest for their role in modulating brain-resident immune cells (microglia) and tumor-associated macrophages (TAMs). Microglia, as key regulators of CNS homeostasis, have been implicated in a range of neurodegenerative disorders, where their dysregulation contributes to disease progression ^1^. Inhibiting CSF1R has shown potential for altering microglial function, reducing neuroinflammation, and promoting neuroprotection ^2,3^. Many preclinical studies use CSF1R inhibitors to temporarily ablate microglia, which resolves disease or injury-induced inflammation, improves brain health and cognitive function ^4-10^.

In neuro-oncology, CSF1R inhibitors are being explored to reprogram TAMs from a tumor-promoting (M2-like) phenotype to a tumor-suppressing (M1-like) phenotype ^11^. Given that TAMs, can comprise up to 50% of the tumor mass, contribute significantly to glioma progression by promoting immune evasion, angiogenesis, and extracellular matrix remodeling ^12,13^, their depletion or reprogramming is a promising therapeutic avenue.

Effective translation of CSF1R inhibitors to CNS indications, however, depends on their ability to reach therapeutic concentrations in the brain. The blood-brain barrier (BBB) serves as a critical physiological defense, tightly regulating the entry of molecules into the central nervous system (CNS). The ability of therapeutic compounds, including CSF1R inhibitors, to effectively penetrate the BBB is crucial to determine their potential efficacy. Despite promising preclinical and early clinical data on the use of CSF1R inhibitors, the potential impact of BBB efflux transporters on the brain bioavailability of these inhibitors remains poorly characterized. Understanding whether pexidartinib and sotuletinib are substrates of the two main drug efflux transporters of the BBB ABCB1 and ABCG2, is crucial in assessing their therapeutic viability for CNS applications.

In addition to limited CNS delivery, CSF1R inhibitors may produce off-target effects, that have been reported on occasionally but are inconsistent between studies and inhibitors. Thus, a better comprehension of the selectivity of CSF1R inhibitors on microglial ablation is warranted, to prevent confounding of interpretation of microglia-targeted interventions.

Here, we systematically evaluate the brain penetration of CSF1R inhibitors by determining their affinity for ABCB1 and ABCG2 *in vitro* and assessing their *in vivo* distribution in murine models. We, next, characterize their effects on microglia, peripheral immune cell subsets, and oligodendrocyte progenitor cells (OPCs), providing a comprehensive assessment of their pharmacological profile relevant to neurodegenerative and neuro-oncology research.

## Methods

### Drugs

Pexidartinib (PLX3397), sotuletinib (BLZ945), and ebvaciclib (PF-06873600) were purchased from MedChemExpress (NJ, USA). Buparlisib (BMK120) was purchased from Syncom (GR, NL). All drugs mentioned above were dissolved in DMSO (Sigma-Aldrich, MO, USA) to a concentration of 10 mM and stored in Eppendorf tubes at −20 °C until use. Elacridar (GF120918) was purchased from GlaxoSmithKline (NC, USA) and zosuquidar (LY335979) from Eli Lilly (IN, USA). Carboxyl-[^14^C]-inulin was generously provided by the Radionuclides Center of the Netherlands Cancer Institute.

### Cell culture

Parental Madin-Darby Canine Kidney (MDCK-parental) cells and sublines transduced with murine *Abcg2* (MDCK-*Abcg2)* or human *ABCG2* (MDCK-*ABCG2*) cDNA and Lilly Laboratories Culture (LLC) porcine kidney epithelial cells (LLC-Pk1) and sublines transduced with murine *Abcb1a* (LLC-*Abcb1a*) or human *ABCB1* (LLC-*ABCB1*) cDNA were cultured in Minimum Essential Medium (MEM) supplemented with 10% fetal bovine serum (FBS), L-glutamine, sodium pyruvate, MEM vitamins, nonessential amino acids, and penicillin/streptomycin (all from Life Technologies, CA, USA) under 37 °C and 5% CO_2_ conditions.

### *In vitro* concentration equilibrium transport assays (CETA)

Cell lines were seeded at a density of 2 × 10^6^ cells per well onto a Transwell^®^ Polycarbonate Membrane (3 µm pores, 24 mm diameter; Costar Corning, NY, USA) in 2 mL complete MEM and cultured for 3 days. After 3 days, all medium in the apical and basal compartment was replaced with complete MEM containing 20% FBS, CSF1R inhibitor cassette (100 nM per drug) and positive control ebvaciclib (100 nM). 15 µL of Carboxyl-[^14^C]-inulin (10^5^ DPM/mL) was added to the basal compartments to assess cellular monolayer integrity (Figure 1A).

**Figure 1.**
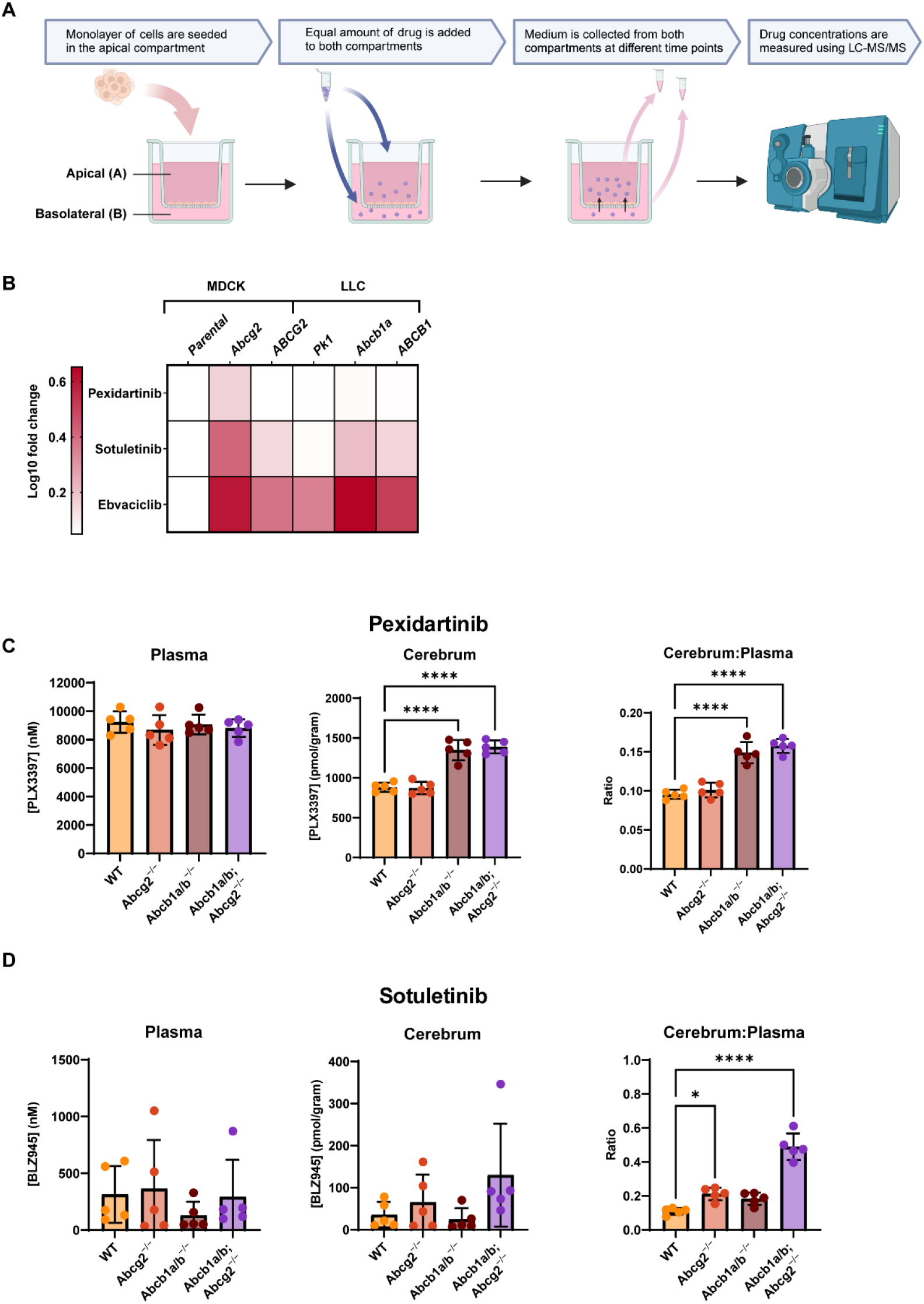
Pharmacokinetic studies using in vitro transwells and in vivo i.v. administration. *(A)* Schematic representation of the experimental set-up of the concentration equilibrium transport assay (CETA). Generated using Biorender O.v.Tellingen.(B) At start, both A and B compartments are loaded with the same concentrations. The heatmap shows the relative gain of drug in the apical versus basal compartment. A darker color implies that the drug is a better substrate of the overexpressed transporter. Ebvaciclib was used as a positive control, being a substrate for both transporters in mice and human. (C-D) Plasma levels of animals receiving a mixture of pexidartinib and sotuletinib were similar across wild type (WT), Abcg2^−/−^, Abcb1a/b^−/−^ and Abcb1a/b^−/−^;Abcg2^−/−^ mice. Note that the within-group variation of sotuletinib is considerably higher than of pexidartinib. A similar higher variation was observed in cerebrum homogenates The cerebrum levels of pexidartinib were significantly higher in Abcb1a/b^−/−^ vs WT mice, but did not further increase in Abcb1a/b^**−*/*−**^;Abcg2^−/−^ mice, indicating that only Abcb1a/b limits the brain penetration. Yet the cerebrum-to-plasma ratio was low across strains for both drugs. Pexidartinib shows increased brain to plasma ratio’s in Abcb1a/b^−/−^ and Abcb1a/b^−/−^;Abcg2^−/−^ mice whereas sotuletinib shows increased brain to plasma ratio’s in Abcg2^−/−^ and a further increase in Abcb1a/b^−/−^;Abcg2^−/−^ mice. * p < 0.05, ** p < 0.01, *** p < 0.001, **** p < 0.0001

Sampling from both compartments was done at 5, 30 minutes, 1, 2 and 4 hours (Figure 1A). Ultima Gold (1:30 v/v) (PerkinElmer, CT, USA), followed by beta-radiation measurement using liquid scintillation counting was used to determine monolayer integrity. Wells were considered leaky and excluded when the Carboxyl-[^14^C]-inulin translocation exceeded 1.5% per hour. The remaining fraction of the aliquots were used for analysis by LC-MS/MS.

The functionality of all transporters was confirmed by adding positive control ebvaciclib, a cyclin-dependent kinase (CDK) inhibitor that is a strong substrate both transporters. Zosuquidar was added at 5 µM to inhibit endogenous porcine ABCB1 in all MDCK cell lines. Elacridar, a dual inhibitor of ABCG2 and ABCB1, was employed to abolish vectorial transport (*i*.*e*., transport only in one direction), thereby controlling whether transport depended solely on those transporters or on the presence of other (unknown) efflux transporters.

For the heatmap representation the 4 hours measurements were used to calculate the Log10(mean apical drug concentration/mean basal drug concentration) for visualization.

### Animals

Male mice were co-housed in Innovive^®^ Individual Ventilated Cages containing bedding material within a temperature-controlled room, following a 12-hour light/dark cycle. They had access to acidified water and food (Transbreed, Technilab-BMI, NB, NL) *ad libitum*. Animal housing and research were approved by the Animal Experimental Committee of the Netherlands Cancer Institute and carried out in accordance with institutional guidelines and national legislation under license AVD3010020198564 (work protocol numbers 21.1.11433 and 26.2.11270).

### *In vivo* brain penetration

The pharmacokinetics of CSF1R inhibitors were first analyzed in wild-type (WT), *Abcg2*^−*/*−^, *Abcb1a/b*^−*/*−^ and *Abcb1a/b-*^*/-*^; *Abcg2*^−*/*−^ FVB mice at 6 to 12 weeks of age (29.6 ± 3.5 grams). For cassette dosing, CSF1R inhibitors were formulated as a mixture in DMSO:Cremophor EL:water (1:1:8 v/v/v) and intravenously injected in the tail vein at a dose of 2.5 mg/kg per drug. Blood and tissues (cerebrum, cerebellum, pons, liver, kidney, and spleen) were harvested and were immediately placed in tubes on ice. Tubes were stored in −20 °C until homogenization.

### *In vivo* PK-PD

We tested repeated oral dosing, in 8-week-old C57BL/6JrJ animals that were treated for 5 consecutive days with either 170 mg/kg/day of sotuletinib dissolved in 20% hydroxypropyl-β-cyclodextrine (HPβCD) (w/v) in H_2_O or 80 mg/kgday of pexidartinib dissolved in 10% DMSO 10% Cremophor 80% H_2_O. Vehicle was 10% DMSO 10% Cremophor in H_2_0 for 3 animals and 20% HPβCD (w/v) in H_2_O for 2 animals. At 2 hours after the final oral gavage blood was collected from the tail vein. Spleens were collected for immediate flow cytometry analysis and brains and organs were isolated and fixed in 10% formalin. From the brain one hemisphere was fresh frozen for drug level measurement.

### Immunohistochemistry staining

Parrafin-embedded brains were sectioned using a microtome (4 µm). For IHC, slides were deparaffinized using xylene. Antigen retrieval was performed using Tris/EDTA pH 9.0 buffer (P2Y12 and F4/80) or Citrate buffer (IBA1). Endogenous peroxidase activity was quenched using 3% H_2_O_2_ in methanol. Slides were pre-incubated with PBS containing 4% BSA and 5% NGS (P2Y12 and F4/80) or 10% milk powder (IBA1) to block nonspecific binding sides. Next, slides were incubated overnight at 4 °C with P2Y12 primary antibody (P2Y12 AS-55043A, Anaspec, CA, USA, 1:100; Iba1 019-19741, Fujifilm WAKO, 1:2000 or F4/80 70076s, Cell Signalling, 1:1000) in PBS containing 1% BSA and 1.25% NGS. After this, slides were incubated with EnVision+ System-HRP Labeled Polymer Anti-Rabbit secondary antibody (K4003; Dako, CA, USA) for 30 minutes. Visualization was achieved using DAB+ substrate (K3468; Dako, CA, USA) for 3 minutes. Counterstaining was performed with hematoxylin for 1 minute. Slides were scanned on a Panoramic P1000 slidescanner.

A quantification method developed by Finney et al. was adapted for use in this project to assess F4/80 DAB coverage. First, stain vectors for color deconvolution were set using the automatic estimation function in Qupath 0.3.2 (NIR, UK). Then the region annotations were defined manually, after which positive pixel identification was done. For DAB+ cell count analysis, four images (each 500 x 500 µm) were made per brain region per mouse and analyzed with QuPath. Cell nuclei were manually counted in each image, and the mean was calculated.

### Flow cytometry analysis of circulating and spleen immune cell populations

Blood of spleen samples were prepared for flow cytometry-analysis as described previously ^14^. In short, blood was first treated with erylysis buffer, blocked with Fc block and then incubated with primary antibody solution in BD Brilliant Buffer and live/dead staining solution (see table 1). The samples were acquired on a 5-laser Aurora full spectrum viewer from Cytek Biosciences. The analysis was done in FlowJo.

**Table 1.**
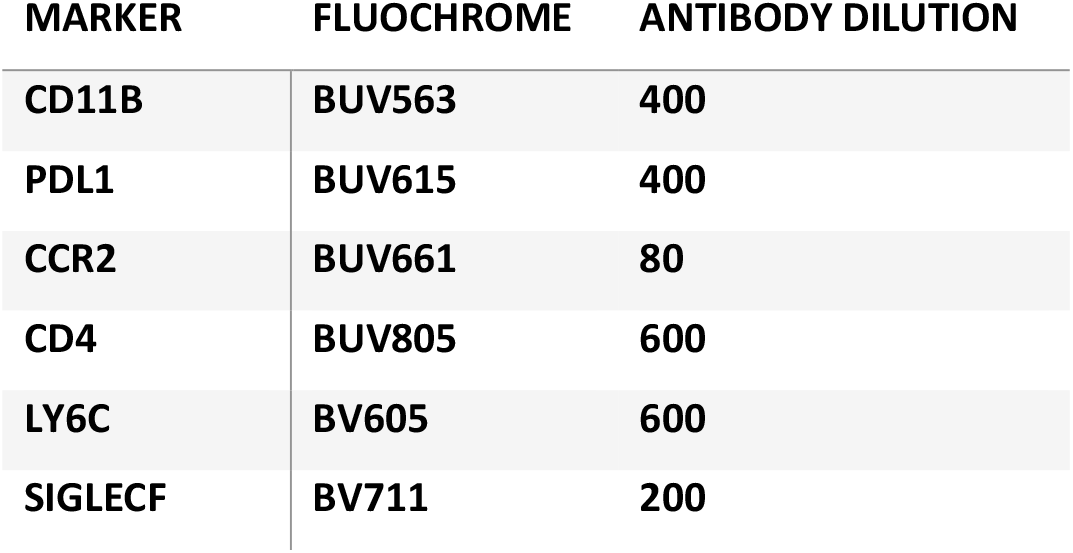
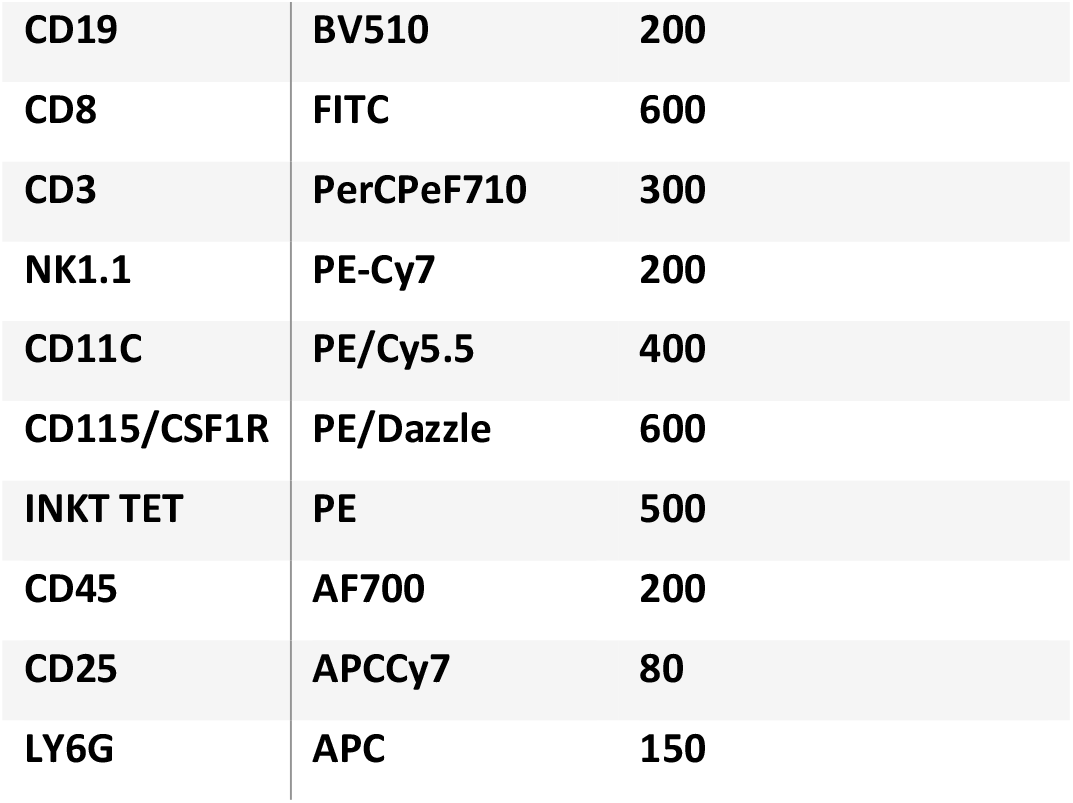
Antibody list for flow cytometry.

| MARKER | FLUOCHROME | ANTIBODY DILUTION |
| --- | --- | --- |
| CD11B | BUV563 | 400 |
| PDL1 | BUV615 | 400 |
| CCR2 | BUV661 | 80 |
| CD4 | BUV805 | 600 |
| LY6C | BV605 | 600 |
| SIGLECF | BV711 | 200 |
| CD19 | BV510 | 200 |
| CD8 | FITC | 600 |
| CD3 | PerCPeF710 | 300 |
| NK1.1 | PE-Cy7 | 200 |
| CD11C | PE/Cy5.5 | 400 |
| CD115/CSF1R | PE/Dazzle | 600 |
| INKT TET | PE | 500 |
| CD45 | AF700 | 200 |
| CD25 | APCCy7 | 80 |
| LY6G | APC | 150 |

### Sample collection and storage

Tissues were weighed and homogenized in 1% Bovine Serum Albumin Fraction V (Roche Diagnostics GmbH, MA, GER) in MiliQ (w/v) using the FastPrep^®^-24 homogenizer (6.0 m/sec, TallPrep, 60 sec; MP-Biomedicals, NY, USA). Plasma was obtained from whole blood by centrifugation (5 min, 5.000 rpm, 4 °C). All tissues and plasma were stored in −20 °C until LC-MS/MS measurement.

### LC-MS/MS analysis

Drug concentrations were measured in samples from the *in vitro* transport assays and *in vivo* pharmacokinetic studies using a liquid chromatography with tandem mass spectrometry (LC-MS/MS). The system was comprised of an UltiMate 3000 LC Systems (Thermo Scientific, Waltham, MA, USA) and a Triple Quad™ 3500 (SCIEX, MA, USA). Data acquisition and quantification were carried out with Analyst 1.7.2 (Sciex) and LC instrument control using Chromeleon 7.2 (Thermo).

### Sample pretreatment

Briefly, calibration standard ranging from 5 nM to 10,000 nM were prepared from a CSF1R inhibitor mixture (100 µM per drug in DMSO) in blank MEM+20%FBS, blank human plasma (Sanquin, Amsterdam, NL) and blank tissue homogenates. Six volumes of internal standard (IS) buparlisib (100 nM in Acetonitrile:Formic acid (99:1 v/v)) was added to one volume of sample while vortex-mixing. Next, samples were centrifuged (5 min, 20.000 g, 4 °C) and the supernatant was diluted in water (1:5 v/v) in a 96 wells plate (Thermo Fisher Scientific, CA, USA). Blanks (IS with blank matrices (MEM, blank human plasma or blank tissue) and double blanks (only blank matrices) were included in each measurement run.

Samples of 50 µL were injected on a Symmetry C18 Column (3.5 µm, 2.1 mm X 150 mm; Waters Corporation, MA, USA) maintained at 50°C. Elution was achieved using a 1.5 minute gradient from 10% to 95% B (mobile phase (A): 0.1% formic acid in water (v/v), mobile phase (B): methanol) with a 0.4 mL/min flow. The 95% B was maintained for 3.5 minutes followed by re-equilibration at 10% B at a 0.6 mL/min flow for 4 min. MS/MS acquisition parameters for the CSF1R inhibitors, buparlisib and ebvaciclib are shown in Table 2.

**Table 2:** Mass spectrometric parameters of pexidartinib, sotuletinib, buparlisib and ebvaciclib. (DP: declustering potential; EP: exit potential; CE: collision energy; CXP: collision cell exit potential)

|  | **Pexidartinib** | **Sotuletinib** | **Buparlisib** | **Ebvaciclib** |
| --- | --- | --- | --- | --- |
| Q1 | 418.2 | 399.1 | 411 | 472.1 |
| Q3 | 258 | 300.9 | 367.1 | 354.1 |
| DP | 130 | 120 | 150 | 90 |
| EP | 10 | 10 | 10 | 10 |
| CE | 37 | 35 | 48 | 43 |
| CXP | 12 | 12 | 20 | 15 |

### Statistical analysis

Data normality was assessed using the Shapiro-Wilk test (α = 0.05). Normally distributed CETA data (Time x Compartment) were analyzed using Repeated-measures two-way ANOVA, based on the General Linear Model. Normally distributed CSF1R inhibitor concentrations in tissues of wild type (WT) were compared with the *Abcg2*^−*/*−^, *Abcb1a/b*^−*/*−^ and *Abcb1a/b; Abcg2*^−*/*−^ FVB mice using a one-way ANOVA, followed by post-hoc Bonferroni tests. Flow cytometry and IHC quantifications were assessed using a one-way ANOVA, followed by post-hoc Bonferroni tests. Statistical significance was considered when *p* < 0.05. All analyses were performed using GraphPad Software version 9.3.0 (MA, USA).

## Results

### *In vitro* transport of CSF1R inhibitors

To determine the affinity of pexidartinib and sotuletinib for the two dominant efflux transporters at the BBB, we used an *in vitro* concentration-equilibrium transwell assay (CETA) using MDCK cells (parental *vs*. those overexpressing (OE) murine Abcg2 or human ABCG2) and LLC cells (parental (PK1) vs murine Abcb1a or human ABCB1) OE cells (Figure 1A). Ebvaciclib was used as positive control and showed strong increases in apical-to-basolateral (A:B) concentrations in all OE cell lines (Figure 1B). Pexidartinib showed only a very modest increase in A:B concentrations in MDCK-Abcg2 OE cells after 2 and 4 hours but not in other cell lines, indicating that pexidartinib is a very weak substrate of Abcg2 *in vitro* and not of ABCG2, Abcb1a or ABCB1 (Figure 1B; S1). Addition of elacridar abolished B to A transport (Figure S1).

Clear B to A transport of sotuletinib was found in the MDCK-Abcg2 OE cell line, whereas transport was more moderate in ABCG2, Abcb1a and ABCB1 OE cells (Figure 1B; S2). Elacridar nearly abolished this transport (Figure S2). These results indicate that sotuletinib is a strong substrate of Abcg2, and a weaker substrate of ABCG2, Abcb1a and ABCB1.

### *In vivo* brain penetration of CSF1R inhibitors

To determine whether the efflux transporters ABCB1a/b or ABCG2 limit the brain accumulation *in vivo*, we used cohorts of wild type (WT), *Abcg2*^−*/*−^, *Abcb1a/b*^−*/*−^ and *Abcb1a/b*^−*/*−^; *Abcg2*^−*/*−^ mice. A mixture of pexidartinib and sotuletinib was administered intravenously at a low dose (2.5 mg/kg of each drug) to minimize animal use while reducing the likelihood of drug-drug interactions. Two hours after dosing, the animals were sacrificed for drug quantification in plasma and tissues. No significant differences in pexidartinib concentrations were observed in plasma, liver, kidney, and spleen levels across strains (Figure S3). Compared with WT mice, pexidartinib levels in the cerebrum, and the corresponding cerebrum-to-plasma ratios were similar in *Abcg2*^*-/-*^ mice but higher in *Abcb1a/b*^−*/*−^ mice (Figure 1C). Relative to *Abcb1a/b*^−*/*−^ mice, levels in *Abcb1a/b*^−*/*−^*;Abcg2*^−*/*−^ mice do not further increase. These results show that Abcb1a, the Abcb1-subtype expressed at the mouse BBB, alone limits the brain penetration of pexidartinib (Figure 1C). Despite higher cerebrum levels in *Abcb1a/b*^−*/*−^ and *Abcb1a/b;Abcg2*^−*/*−^ mice, cerebrum-to-plasma ratios remained very low (0.09-0.16)(Figure 1C). Notably, tissue-to-plasma ratio in liver, kidney and spleen were also relatively low (i.e. 0.3 to 0.5), indicating that pexidartinib is not extensively distributed into peripheral tissues.

The brain accumulation of sotuletinib is limited by both Abcg2 and Abcb1a (Figure 1D). Notably, the within-group variabilities in sotuletinib levels in plasma were considerably higher than that of pexidartinib. This was not due to dosing issues, as both drugs were administered together. Similar variabilities were seen in all tissues, but this was largely corrected by using tissue-to-plasma ratios (Figure S3). The cerebrum-to-plasma ratio in WT mice is 0.1 and increases by 2-fold in *Abcg2*^*-/-*^ and *Abcb1a/b*^*-/-*^ and by 5-fold *Abcb1a/b*^−*/*−^*;Abcg2*^−*/*−^ mice, respectively (Figure 1D). The accumulation of sotuletinib is approximately 1.7 in liver, 1.2 in kidney and 0.7 in spleen across mouse strains (Figure S3). The relative brain distribution of sotuletinib in WT mice (0.1) compared to other tissues (e.g. liver: 1.7) is lower for sotuletinib, than for pexidartinib (Figure 1C-D; S3).

### Five-day CSF1R inhibitor treatment affects CSF1R expressing monocytes in blood and spleen

To determine the effects of CSF1R inhibitor treatment on tissue resident macrophages and immune populations, we subjected animals to dose levels that, based on literature, is expected to result in rapid microglial depletion. For sotuletinib, 169mg/kg by oral gavage was previously used ^15,18^, while for pexidartinib 600 ppm dosing via the food was reported^16,19-21^. The latter is estimated to correspond to approximately 80mg/kg/day (based on 4g food intake with a mouse body weight of 30). We treated mice with 80mg/kg pexidartinib or 170 mg/kg sotuletinib, both given p.o. daily for 5 consecutive days. Immune cell phenotyping was performed 24 hours after the last dose (Figure 2A). Plasma levels at 2 hours after oral drug administration were extremely high drug (2,860 µM for pexidartinib and 1,340 µM for sotuletinib)(Figure 2B). At sacrifice (24 h post administration), the plasma levels were reduced to about 6.5 and 7.6 of µM of pexidartinib and sotuletinib, respectively, implicating plasma half-lives of approximately 2.5 and 3 h, respectively. The cerebrum-to-plasma ratio of pexidartinib was again low, being 0.15 and 0.4 for pexidartinib and sotuletinib, respectively, (Table 3; Figure 2B).

**Table 3.** Available mouse pharmacokinetics on pexidartinib and sotuletinib.

| **Drug** | **First author, year** | **Dose** | **Timing Pk measurement** | **Plasma levels (nM)** | **Brain levels (nM)** | **Brain-to-plasma ratio** |
| --- | --- | --- | --- | --- | --- | --- |
| Sotulentinib | Beckman 2018<sup>15</sup> | 169mg/kg, oral gavage | not reported | ± 70000 | ± 35000 | 0.58 |
| **Sotulentinib** | **This study** | **2.5mg/kg intravenous** | **2h** | **313** | **35** | **0.1** |
| **Sotulentinib** | **This study** | **170mg/kg, oral gavage** | **2h** | **1340000** |  |  |
| **Sotulentinib** | **This study** | **170mg/kg, oral gavage** | **24h** | **7610** | **3060** | **0.4** |
| Pexidartinib | Najafi 2018<sup>16</sup> | 660mg/kg in diet, estimated intake 88mg/kg | continous exposure | 160000 | 8 | 0.05 |
| Pexidartinib | Elmore 2014<sup>17</sup> | 290mg/kg in diet, estimated intake 39mg/kg | continous exposure | 140000 | 5 | 0.03 |
| **Pexidartinib** | **This study** | **2.5mg/kg intravenous** | **2h** | **9240** | **880** | **0.1** |
| **Pexidartinib** | **This study** | **80mg/kg oral gavage** | **2h** | **2860000** |  |  |
| **Pexidartinib** | **This study** | **80mg/kg oral gavage** | **24h** | **6496** | **934** | **0.15** |

**Figure 2.**
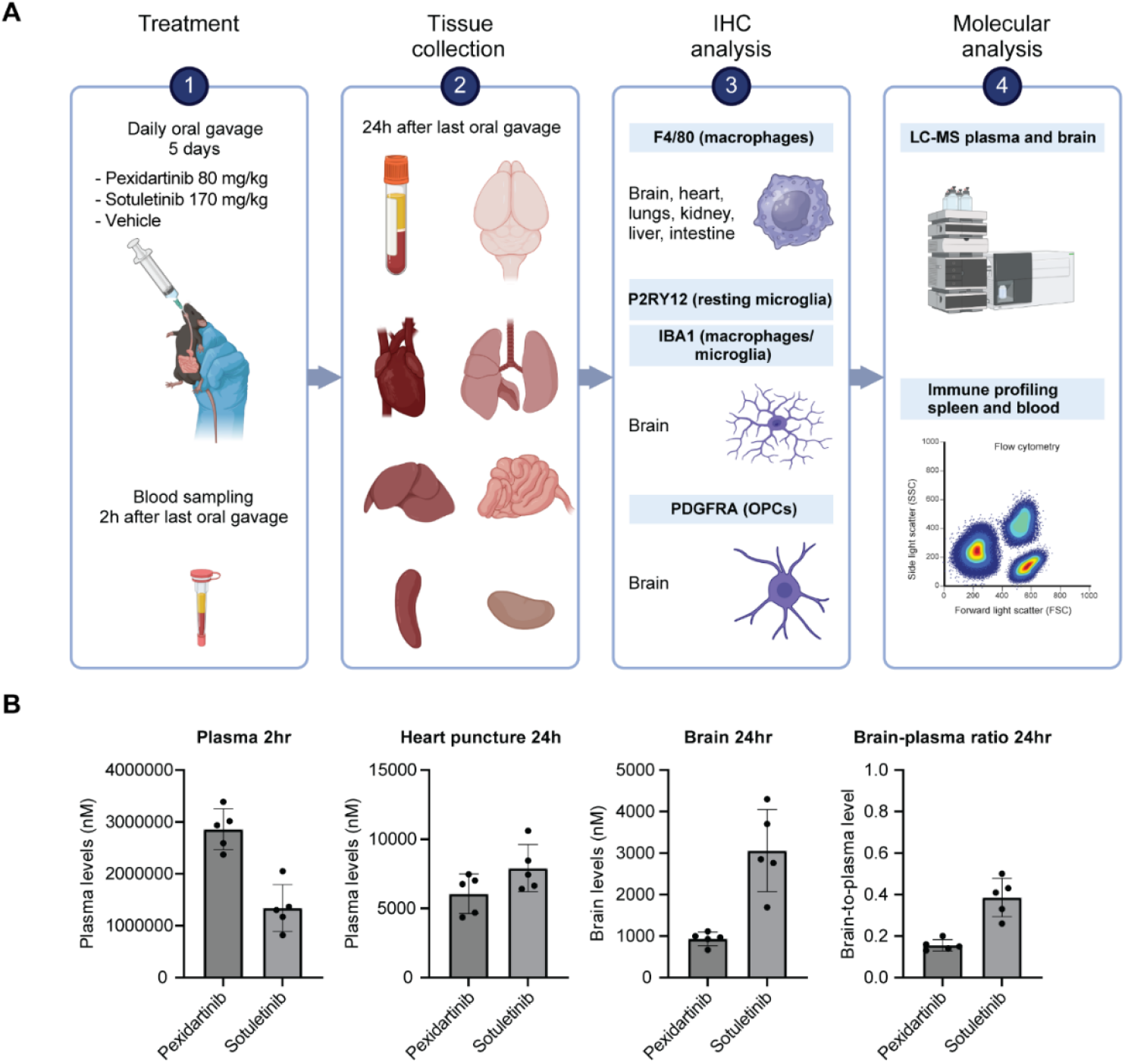
Pharmacodynamic study assessing the effects of 5-day Csf1r inhibitor treatment on immune populations. A. Schematic overview of the experimental set-up where animals were treated orally with either pexidartinib, sotuletinib or vehicle for 5 consecutive days. Organs were collected for IHC, plasma and brain drug levels were measured by LC-MS/MS. Blood and spleen immune-profiling was performed by flow cytometry. Generated using Biorender O.v.Tellingen. B. Drug levels in plasma from blood collected from the tail vein at 2 h and at sacrifice (24 h) by heart puncture, together with brain samples. Brain to plasma ratios were calculated from the 24h samples.

Flow cytometry analysis of immune populations in blood shows selective ablation of Ly6C-monocytes and CSF1R+ monocytes upon treatment with either inhibitor. In the spleen only pexidartinib resulted in ablation of Ly6C-monocytes. Both drugs ablated CSF1R+ monocytes in the spleen. On the other hand, pexidartinib treatment modestly induced CD4+ and Cd8+ T cells while reducing dendritic cells in the spleen. In the blood, neutrophils were induced whereas Cd8+ T-cells were decreased upon pexidartinib treatment (Figure 3).

**Figure 3.**
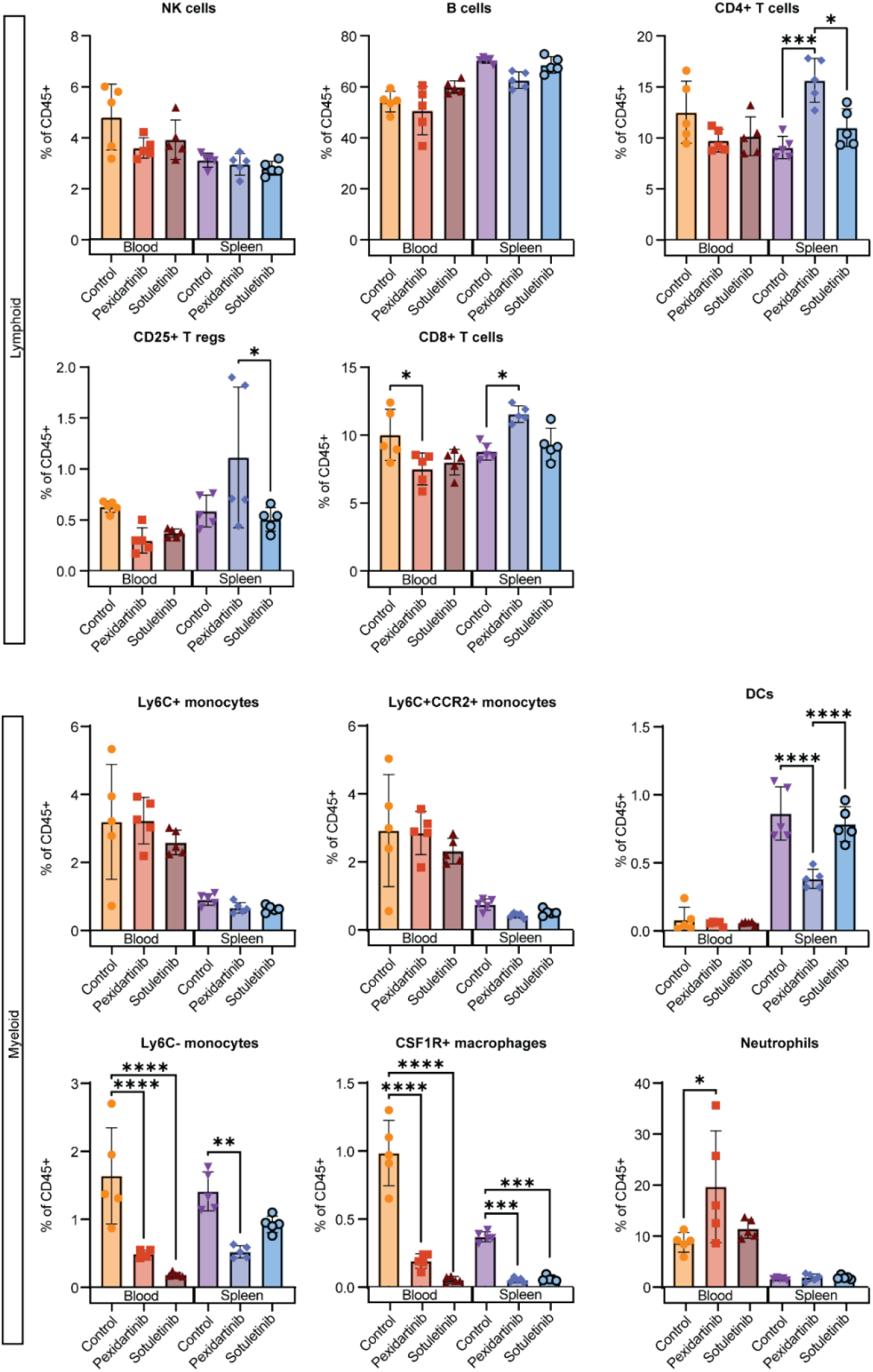
Flow cytometry-analysis of immune cells on blood and spleen 24 hours after the last oral gavage. Analysis of 5 populations of lymphoid cells and 7 populations of myeloid cells was performed showing some drug specific effects such as pexidartinib increases CD4+ and CD8+ T-cells in the spleen and reduces CD8+ T-cells in blood. Pexidartinib reduces dendritic cells (DCs) and Ly6C-monocytes in the spleen. Pexidartinib induced neutrophil levels in blood. Ly6C-monocytes were reduced in the blood upon both inhibitors as well as reduction of CD115+ (CSF1R) macrophages in blood and spleen. * p < 0.05, ** p < 0.01, *** p < 0.001, **** p < 0.0001

### CSF1R inhibitors reduce macrophage coverage in liver, intestine, kidney and brain

To determine the effects of CSF1R inhibitor treatment on tissue (resident) macrophages in organs, IHC for the pan-macrophage marker F4/80 was performed. We observed a strong reduction in macrophage coverage in the kidney and more modestly in intestine and liver (Figure 4). In general, sotuletinib had more impact than pexidartinib at these dose levels (Figure 4). The variabilities in the heart make results in this tissue inconclusive (Figure 4).

**Figure 4.**
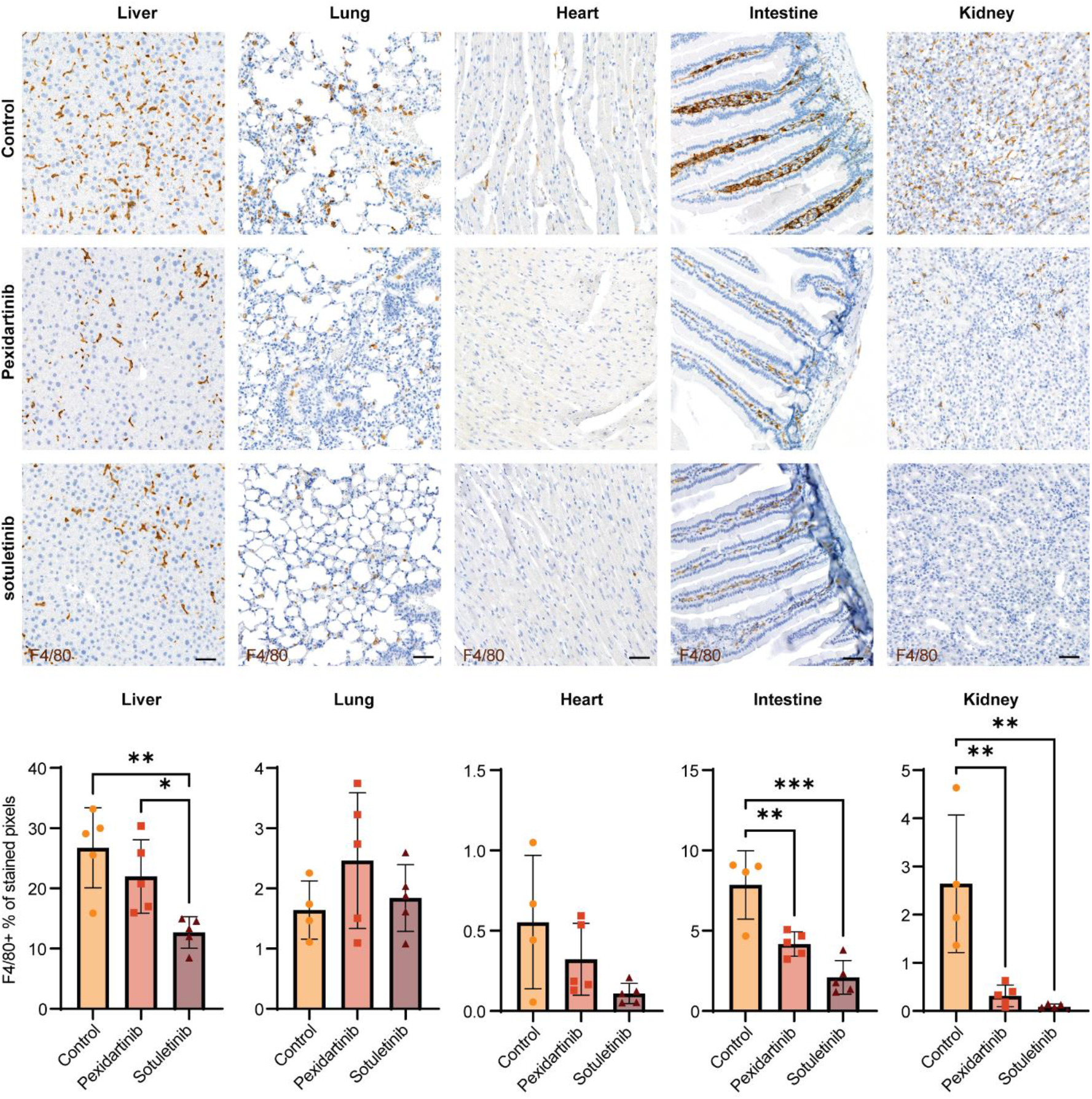
IHC analysis of macrophages in various organs shows organ-specific effects of CSF1R inhibitors. A. Both inhibitors reduced macrophage coverage in intestine and kidney, whereas only sotuletinib reduced coverage in the liver (in line with the high levels of sotuletinib found in the liver in the Pk experiment). In the heart macrophages also seem affected, however due to the high variability observed in the control group these differences were not statistically significant. B. Macrophage coverage (F4/80+% of stained pixels) and numbers (counts) were reduced upon both inhibitors in all brain regions assessed. Regarding coverage there is a significant difference between the two inhibitors with pexidartinib inducing a milder effect compared to sotuletinib. Scale bar represents 50 micron. * p < 0.05, ** p < 0.01, *** p < 0.001

In the brain, F4/80 cells (microglia and CNS macrophages) are depleted upon treatment with either inhibitor (Figure 5A; S4A). Microglial depletion was validated using two additional markers: P2RY12, a marker labelling homeostatic microglia but not reactive microglia, monocytes and perivascular macrophages, and Iba1, a pan-macrophage marker that is most widely used in literature to stain microglia and perivascular macrophages in the brain regardless of reactivity of microglia. Compared to other tissues, except kidney, we found a very profound 85-88% reduction of microglia in the brain with both inhibitors (Figure 5B-C; S4B-C). We also observed morphological changes in the remaining microglia (Figure 5A-C; S4A-C).

**Figure 5.**
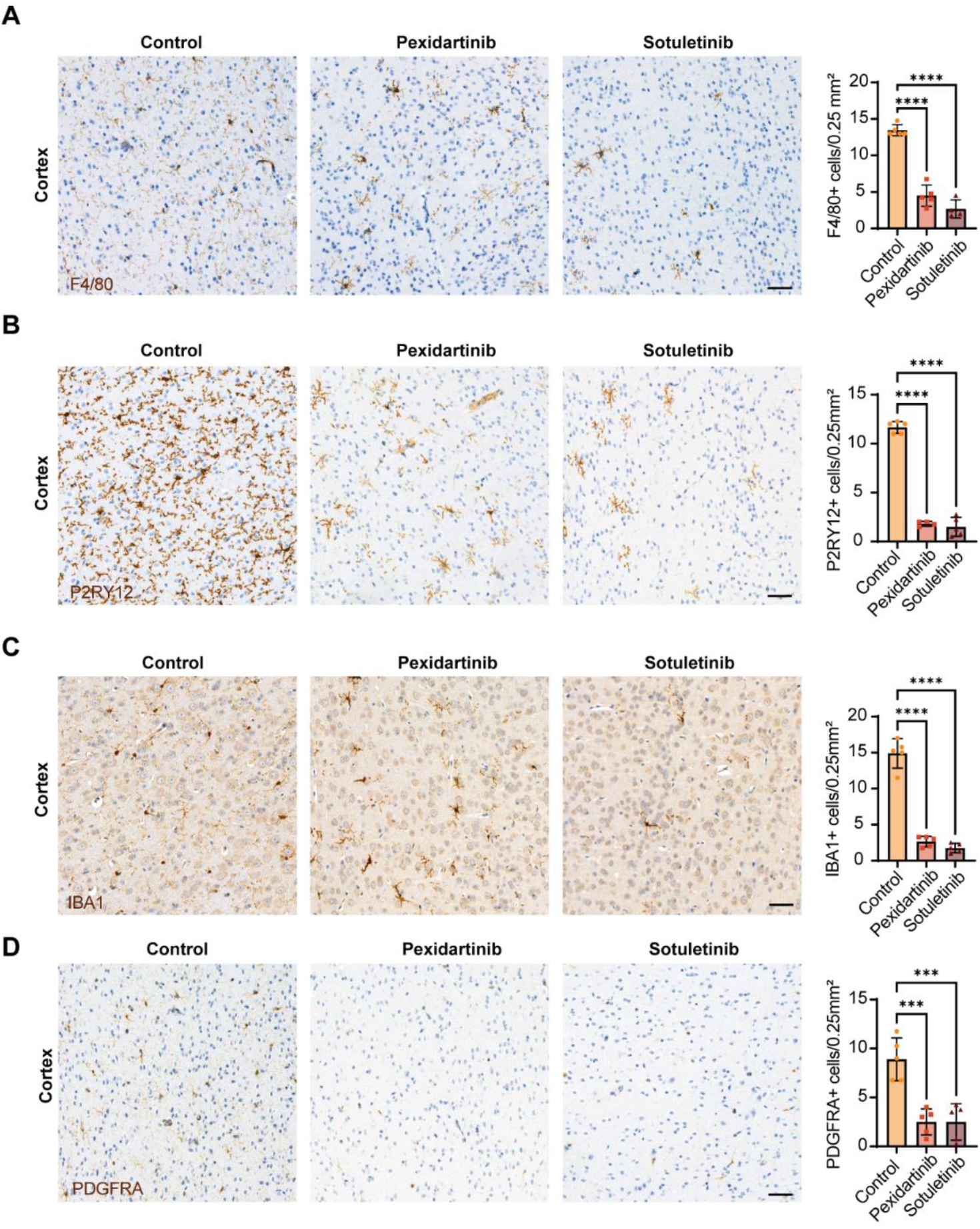
Assessment of microglia and oligodendrocyte progenitor cells in the cortex. Both inhibitors resulted in comparable microglial ablation levels measured by the pan-macrophage markers F4/80 (A) and IBA1 (C) as well as the homeostatic microglial specific marker P2RY12 (B). Both inhibitors show significant reductions of oligodendrocyte progenitors cells using PDGFRA (D). Scale bar represents 50 micron. * p < 0.05, ** p < 0.01, *** p < 0.001

Besides microglia, we also checked the effects of the CSF1R inhibitors on oligodendrocyte progenitor cells (OPCs) as they express PDGFRA, which is closely related to CSF1R. Notably, we also observed a 78% reduction of OPCs (Figure 5D; S4D), indicating that these inhibitors can also target this brain cell population.

## Discussion

Although CSF1R inhibitors are already used in clinical trials for glioblastoma (NCT02829723, NCT01790503, NCT01349036, NCT02452424) and amyotrophic lateral sclerosis (NCT04066244), there is very limited information about their ability to cross the blood-brain barrier (BBB) and/or the influence of the drug efflux transporters ABCB1 and ABCG2 that are expressed at the BBB. Therefore, we used *in vitro* and *in vivo* models to investigate the affinity of pexidartinib and sotuletinib for the efflux transporters ABCG2 and ABCB1, and their impact on the brain penetration. *In vitro*, sotuletinib was a weak substrate for mouse and human ABCB1 and a strong substrate for mouse ABCG2. Pexidartinib was only a weak substrate for mouse Abcg2. The *in vivo* studies showed that sotuletinib brain distribution is limited by both Abcb1 and Abcg, whereas pexidartinib is restricted primarily by Abcb1. The brain distribution for both compounds was low. Brain-to-plasma (B/P) ratios of pexidartinib increased only marginally from 0.10 in wild-type mice to 0.15 in transporter-deficient strains, while the B/P ratio of sotuletinib increased to about 0.4. Compounds with low affinity for ABCB1 and or ABCG2 often exhibit reasonable brain accumulation. The poor accumulation of pexidartinib in brain was accompanied by a relatively low accumulation in other tissues as well and implicates a low volume of distribution. These values align with previous reports of limited CNS exposure for pexidartinib and sotuletinib (Table 3).

We next assessed the systemic and CNS effects of repeated high-dose oral dosing. Both inhibitors efficiently depleted microglia (pexidartinib: 82%, sotuletinib: 88%) despite low brain penetration, indicating that microglia may be particularly sensitive to CSF1R blockade. We found efficient depletion of tissue (resident) macrophages and ly6C-monocytes in blood upon either CSF1R inhibitor treatment, in line with previous reports ^22,23^. The effects of sotuletinib in other tissues seems stronger compared to pexidartinib, although these differences are not always statistically significant. Ly6C^+^ monocytes, which are critical for repopulating tissue-resident macrophages, were largely unaffected by either inhibitor—consistent with some reports^21^ but contrasting with others^23^. The increase in CD4+ T cells in the spleen upon pexidartinib treatment in the current study is opposite from the findings of a previous study ^23^. Whereas, the reductions of circulating CD8+ T cells observed upon pexidartinib treatment confirm a previous study ^19^. We confirm ablation of liver macrophages by either inhibitor ^19,22^. Notably, we demonstrate that CSF1R inhibition also affects, although to a lesser extend compared to microglia, macrophage populations in spleen, heart, intestine, and kidney—tissues not extensively studied in this context.

A key concern emerging from our work is the profound ablation of OPCs by these high-dose CSF1R inhibitor treatment required for microglial ablation. Although these inhibitors do not target PDGFRA, they might inhibit other tyrosine kinases (e.g. C-KIT, FLT3 and PDGFRB^24,25^), due to structural similarity. OPCs are known to be dependent on PDGFRA signaling for their survival ^26^. Others have also reported on reductions on OPC numbers upon pexidartinib ^27^, sotuletinib ^15,18^ and even the more selective CSF1R inhibitor PLX5622^27,28^ treatment *in vivo*. OPC depletion has the potential to confound interpretations in studies aiming to improve brain health or cognition. OPCs likely play an important role in neurodegenerative diseases given their involvement in myelin repair, including diseases for which CSF1R inhibition has been studied as an intervention ^29,30^. Moreover, OPC loss itself can impair cognitive performance^31^. Thus, OPC depletion represents a potential adverse consequence of CSF1R-targeted therapies

Prior work has shown variable effects on OPC survival depending on the CSF1R inhibitor used, the dose, and the route of administration ^16,27^. For instance, continuous dietary or drinking-water dosing generally results in lower peak exposures and may produce milder OPC loss, whereas bolus oral gavage, as used here, likely produces high peak concentrations that exacerbate off-target effects. Future studies should investigate whether continuous lower-level exposure can maintain efficient microglial ablation while sparing OPCs, particularly for sotuletinib, for which dietary administration has not yet been evaluated. Such approaches may help preserve the therapeutic potential of CSF1R inhibition while minimizing unintended disruption to other essential glial populations.

## Conclusion

In summary, this study provides further insight into the brain penetration of CSF1R inhibitors and the role of drug transporters. We demonstrate that pexidartinib and sotuletinib are not only retained by efflux proteins, but exert generally low brain penetrability, even in the absence of transporters. Nevertheless, both inhibitors show excellent ablative capacities of microglia and macrophages in various organs, non-classical monocytes in blood, with limited effects on other immune populations in blood and spleen. However, of concern is the apparent loss of OPCs upon CSF1R inhibitor treatment at relatively high doses to acquire rapid microglia ablation. Furthermore, these effects are established at very high drug exposure levels that are most likely not clinically achievable. Future studies should investigate whether positive effects can be obtained using clinically relevant concentrations and delivery routes suitable for optimal on-target effects and limited effects on OPCs.

## References

1 Gao, C., Jiang, J., Tan, Y. & Chen, S. Microglia in neurodegenerative diseases: mechanism and potential therapeutic targets. Signal Transduct Target Ther 8, 359, doi:10.1038/s41392-023-01588-0 (2023).

2 Han, J. et al. Inhibition of colony stimulating factor-1 receptor (CSF-1R) as a potential therapeutic strategy for neurodegenerative diseases: opportunities and challenges. Cell Mol Life Sci 79, 219, doi:10.1007/s00018-022-04225-1 (2022).

3 AmeliMojarad, M. & AmeliMojarad, M. The neuroinflammatory role of microglia in Alzheimer’s disease and their associated therapeutic targets. CNS Neurosci Ther 30, e14856, doi:10.1111/cns.14856 (2024).

4 Gibson, E. M. et al. Methotrexate Chemotherapy Induces Persistent Tri-glial Dysregulation that Underlies Chemotherapy-Related Cognitive Impairment. Cell 176, 43–55 e13, doi:10.1016/j.cell.2018.10.049 (2019).

5 Acharya, M. M. et al. Elimination of microglia improves cognitive function following cranial irradiation. Sci Rep 6, 31545, doi:10.1038/srep31545 (2016).

6 Henry, R. J. et al. Microglial Depletion with CSF1R Inhibitor During Chronic Phase of Experimental Traumatic Brain Injury Reduces Neurodegeneration and Neurological Deficits. J Neurosci 40, 2960–2974, doi:10.1523/JNEUROSCI.2402-19.2020 (2020).

7 Liu, H. et al. Microglial repopulation alleviates age-related decline of stable wakefulness in mice. Front Aging Neurosci 14, 988166, doi:10.3389/fnagi.2022.988166 (2022).

8 Ritzel, R. M. et al. Brain injury accelerates the onset of a reversible age-related microglial phenotype associated with inflammatory neurodegeneration. Sci Adv 9, eadd1101, doi:10.1126/sciadv.add1101 (2023).

9 Iba, M. et al. Microglial and neuronal fates following inhibition of CSF-1R in synucleinopathy mouse model. Brain Behav Immun 123, 254–269, doi:10.1016/j.bbi.2024.09.016 (2025).

10 Cheng, X. et al. Repopulated retinal microglia promote Muller glia reprogramming and preserve visual function in retinal degenerative mice. Theranostics 13, 1698–1715, doi:10.7150/thno.79538 (2023).

11 Wang, G. et al. Tumor-associated microglia and macrophages in glioblastoma: From basic insights to therapeutic opportunities. Front Immunol 13, 964898, doi:10.3389/fimmu.2022.964898 (2022).

12 Soto, M. S. & Sibson, N. R. The Multifarious Role of Microglia in Brain Metastasis. Front Cell Neurosci 12, 414, doi:10.3389/fncel.2018.00414 (2018).

13 Wen, J., Wang, S., Guo, R. & Liu, D. CSF1R inhibitors are emerging immunotherapeutic drugs for cancer treatment. Eur J Med Chem 245, 114884, doi:10.1016/j.ejmech.2022.114884 (2023).

14 Taranto, D. et al. Multiparametric Analyses of Hepatocellular Carcinoma Somatic Mouse Models and Their Associated Tumor Microenvironment. Curr Protoc 1, e147, doi:10.1002/cpz1.147 (2021).

15 Beckmann, N. et al. Brain region-specific enhancement of remyelination and prevention of demyelination by the CSF1R kinase inhibitor BLZ945. Acta Neuropathol Commun 6, 9, doi:10.1186/s40478-018-0510-8 (2018).

16 Najafi, A. R. et al. A limited capacity for microglial repopulation in the adult brain. Glia 66, 2385–2396, doi:10.1002/glia.23477 (2018).

17 Elmore, M. R. et al. Colony-stimulating factor 1 receptor signaling is necessary for microglia viability, unmasking a microglia progenitor cell in the adult brain. Neuron 82, 380–397, doi:10.1016/j.neuron.2014.02.040 (2014).

18 Hagemeyer, N. et al. Microglia contribute to normal myelinogenesis and to oligodendrocyte progenitor maintenance during adulthood. Acta Neuropathol 134, 441–458, doi:10.1007/s00401-017-1747-1 (2017).

19 Claeys, W. et al. Limitations of PLX3397 as a microglial investigational tool: peripheral and off-target effects dictate the response to inflammation. Front Immunol 14, 1283711, doi:10.3389/fimmu.2023.1283711 (2023).

20 Hohsfield, L. A. et al. Subventricular zone/white matter microglia reconstitute the empty adult microglial niche in a dynamic wave. Elife 10, doi:10.7554/eLife.66738 (2021).

21 Okojie, A. K. et al. Distinguishing the effects of systemic CSF1R inhibition by PLX3397 on microglia and peripheral immune cells. J Neuroinflammation 20, 242, doi:10.1186/s12974-023-02924-5 (2023).

22 Pognan, F. et al. Liver enzyme delayed clearance in rat treated by CSF1 receptor specific antagonist Sotuletinib. Curr Res Toxicol 3, 100091, doi:10.1016/j.crtox.2022.100091 (2022).

23 Han, J. et al. Underestimated Peripheral Effects Following Pharmacological and Conditional Genetic Microglial Depletion. Int J Mol Sci 21, doi:10.3390/ijms21228603 (2020).

24 Pyonteck, S. M. et al. CSF-1R inhibition alters macrophage polarization and blocks glioma progression. Nat Med 19, 1264–1272, doi:10.1038/nm.3337 (2013).

25 Voissiere, A. et al. The CSF-1R inhibitor pexidartinib affects FLT3-dependent DC differentiation and may antagonize durvalumab effect in patients with advanced cancers. Sci Transl Med 16, eadd1834, doi:10.1126/scitranslmed.add1834 (2024).

26 Dang, T. C. et al. Powerful Homeostatic Control of Oligodendroglial Lineage by PDGFRalpha in Adult Brain. Cell Rep 27, 1073–1089 e1075, doi:10.1016/j.celrep.2019.03.084 (2019).

27 Liu, Y. et al. Concentration-dependent effects of CSF1R inhibitors on oligodendrocyte progenitor cells ex vivo and in vivo. Exp Neurol 318, 32–41, doi:10.1016/j.expneurol.2019.04.011 (2019).

28 Dagher, N. N. et al. Colony-stimulating factor 1 receptor inhibition prevents microglial plaque association and improves cognition in 3xTg-AD mice. J Neuroinflammation 12, 139, doi:10.1186/s12974-015-0366-9 (2015).

29 Clayton, B. L. L. & Tesar, P. J. Oligodendrocyte progenitor cell fate and function in development and disease. Curr Opin Cell Biol 73, 35–40, doi:10.1016/j.ceb.2021.05.003 (2021).

30 Zou, P., Wu, C., Liu, T. C., Duan, R. & Yang, L. Oligodendrocyte progenitor cells in Alzheimer’s disease: from physiology to pathology. Transl Neurodegener 12, 52, doi:10.1186/s40035-023-00385-7 (2023).

31 Brousse, B. et al. Characterization of a new mouse line triggering transient oligodendrocyte progenitor depletion. Sci Rep 13, 21959, doi:10.1038/s41598-023-48926-4 (2023).

