## Supplemental figures for "CSF1R inhibitors pexidartinib and sotuletinib induce rapid glial ablation despite their limited brain penetrability"

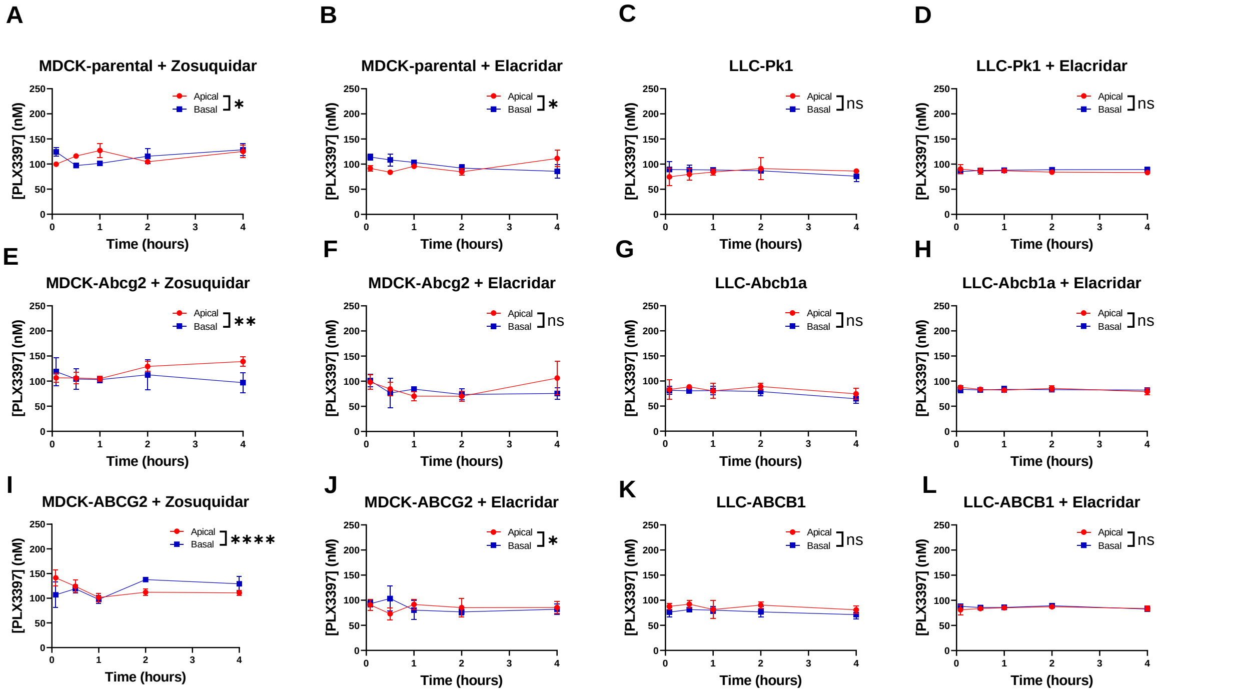


**Figure S1** ***In vitro*** **transport of pexidartinib using concentration equilibrium transport assays** conducted with MDCK or LLC cells expressing murine *Abcg2* and human *ABCG2,* and murine *Abcb1a* and human *ABCB1*, respectively. No transport mediated by *Abcb1/ABCB1* was observed for pexidartinib, while minimal transport was detected in the MDCK-Abcg2 cell lines. Data are presented as mean ± SD (n = 2-4); ns: not significant, *: $p < 0.05$, **: $p < 0.01$, ****: $p < 0.0001$.

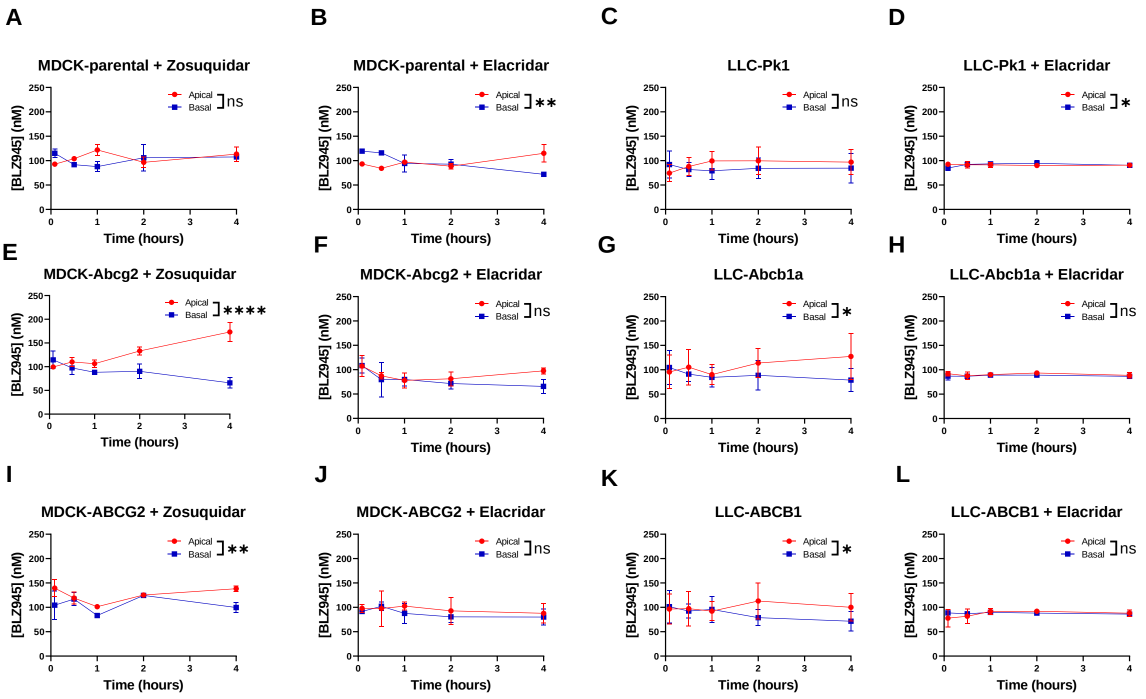


**Figure S2** ***In vitro*** **transport of sotuletinib using concentration equilibrium transport assays** conducted with MDCK or LLC cells expressing murine *Abcg2* and human *ABCG2*, and murine *Abcb1* and human *ABCB1*, respectively. Significant transport mediated by *Abcg2 a*nd *ABCG2* was observed for sotuletinib, with transport also detected in the LLC cell lines. This transport was eliminated upon the addition of Elacridar. Data are presented as mean ± SD (n = 2-4); ns: not significant, *: $p < 0.05$, **: $p < 0.01$, ****: $p < 0.0001$.

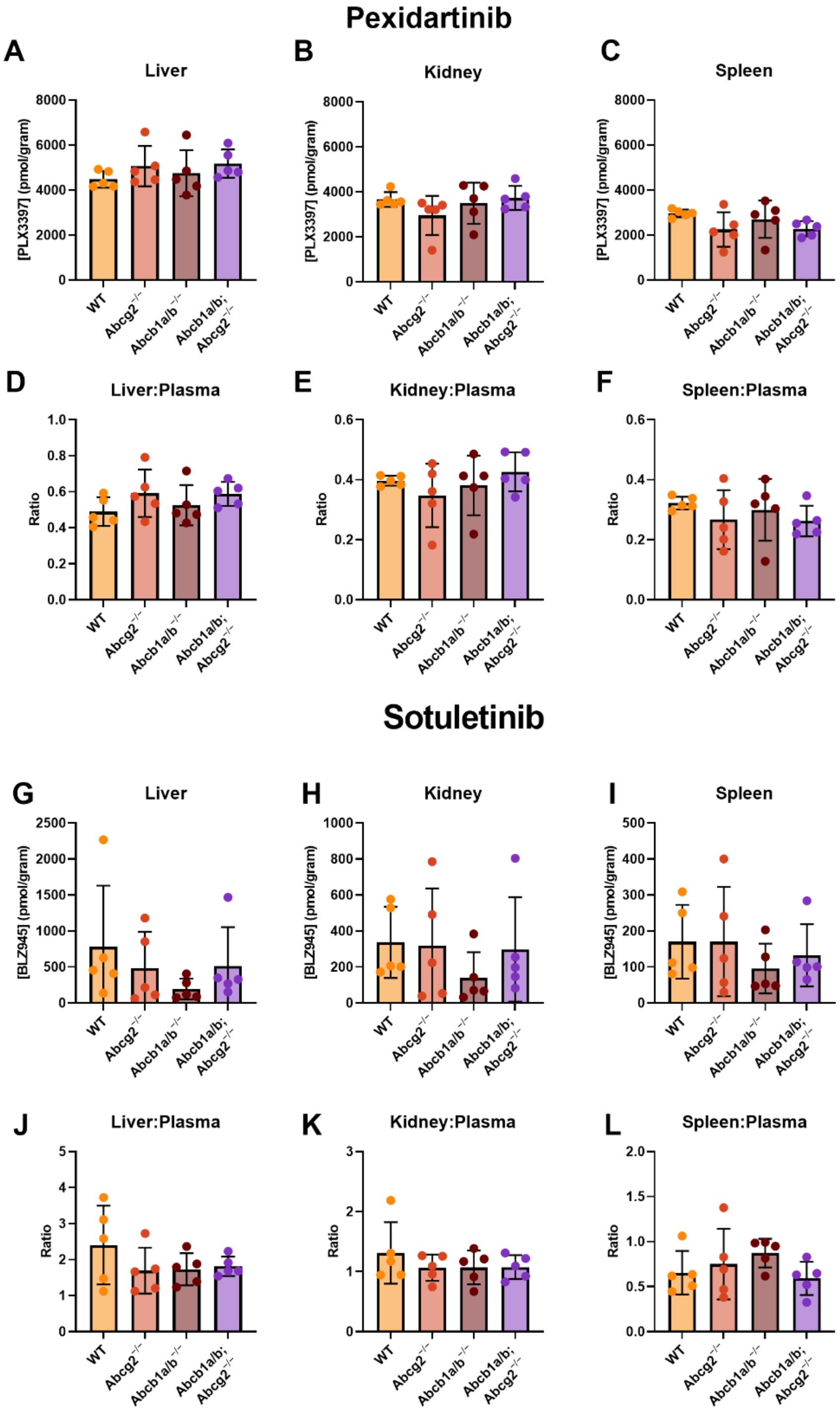


***Figure S3. Plasma and organ drug levels of pexidartinib and sotuletinib 2 hours after i.v. administration.*** *Levels in all organs and ratio's between organs versus plasma were similar amongst the wild type (WT), Abcg2$^{-/-}$, Abcb1a/b$^{-/-}$ and Abcb1a/b$^{-/-}$;Abcg2$^{-/-}$ mice.*

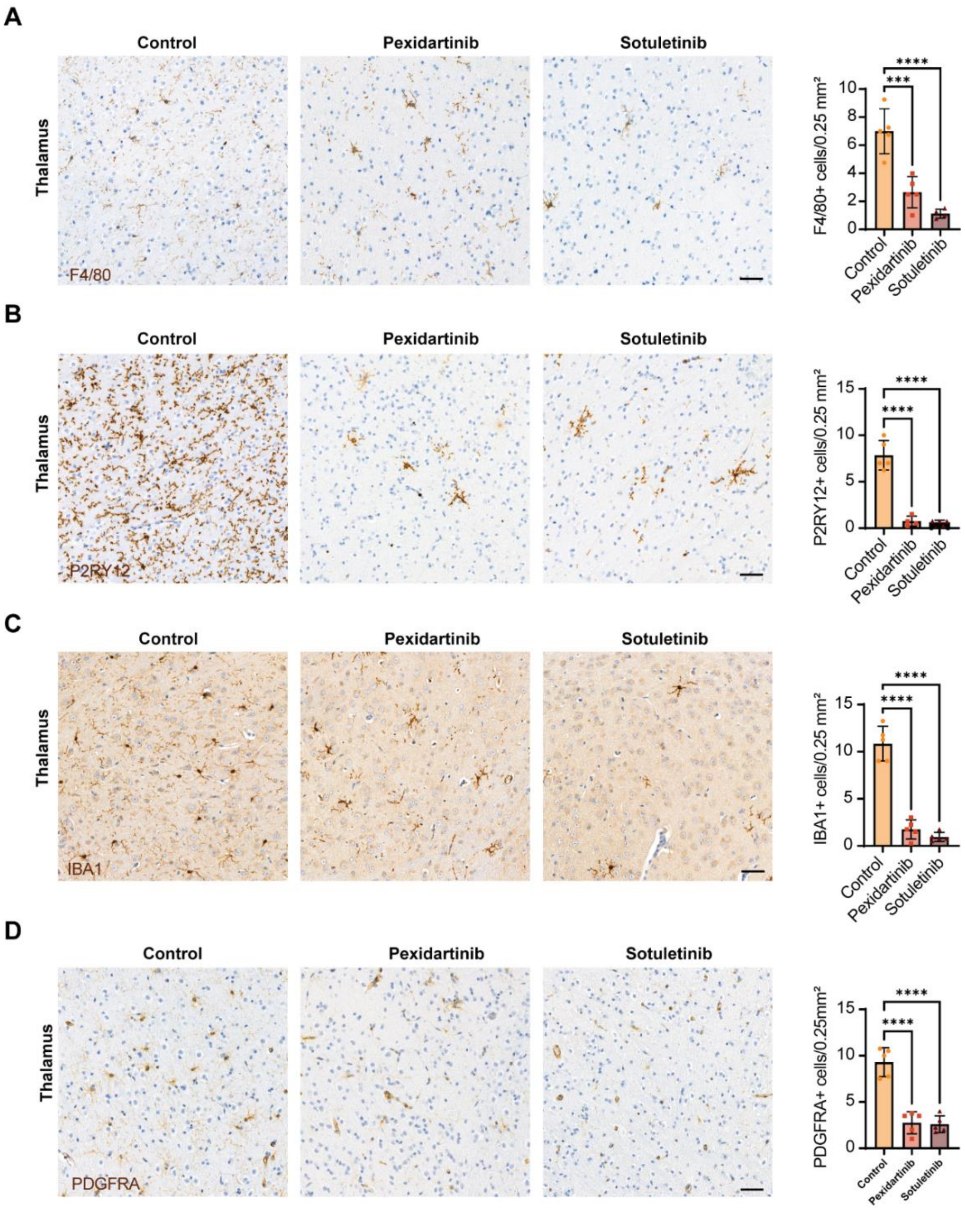


***Figure S4.*** **Assessment of microglia and oligodendrocyte progenitor cells in the thalamus**

Both inhibitors resulted in comparable microglial ablation levels measured by the pan-macrophage markers F4/80 (A) and IBA1 (C) as well as the homeostatic microglial specific marker P2RY12 (B). Both inhibitors show significant reductions of oligodendrocyte progenitors cells using PDGFRA (D). Scale bar represents 50 micron. * $p < 0.05$, ** $p < 0.01$, *** $p < 0.001$, **** $p < 0.0001$
